# How a Disease-Associated Mutation Destabilizes the Sliding Clamp PCNA

**DOI:** 10.64898/2026.11.15.766507

**Authors:** Joshua Pajak, Kendra E. Baker, Joseph Magrino, Brian A. Kelch

## Abstract

The eukaryotic sliding clamp, Proliferating Cell Nuclear Antigen (PCNA), is a homo-trimeric ring-shaped protein complex that encircles DNA and serves as a scaffold for enzymes during DNA replication and the DNA damage response. Patients harboring the homozygous point mutation C148S in PCNA have the genetic disease <u>P</u>CNA <u>A</u>ssociated DNA <u>R</u>epair <u>D</u>isorder (PARD). Our lab showed that this PCNA variant has a drastic 10 °C thermostability defect. However, the crystal structure of the C148S variant displays a nearly identical conformation as WT PCNA, so it remained unclear how this point mutation destabilizes PCNA. Here, we use extensive all-atom classical molecular dynamics and well-tempered metadynamics simulations to predict how this mutation destabilizes PCNA. We first show that the C148S substitution changes an internal hydrogen-bond network and permits water molecules to transiently bind within the protein core, distorting the α-helix containing C148S. This distortion in turn weakens the subunit interface, providing a molecular explanation for the lowered stability of the PCNA ring. Thus, our study addresses the molecular mechanism by which this disease-associated variant destabilizes PCNA and highlights the importance of understanding protein dynamics to interpret the effects of mutations in disease.

## Introduction

All life uses sliding clamps to help replicate and repair DNA. <u>P</u>roliferating <u>C</u>ell <u>N</u>uclear <u>A</u>ntigen (PCNA) is the eukaryotic sliding clamp protein complex [1,2]. PCNA forms a homo-trimeric ring that encircles DNA, and coordinates over two-hundred binding partners that regulate and drive DNA metabolism. It is critical that chromatin-bound PCNA levels are maintained at the correct amounts and with correct timings, as misregulation of PCNA levels on chromatin leads to genome-wide instabilities [3–6], and exhaustion of PCNA regulates the S-phase checkpoint [7].

Because PCNA is a closed ring in solution [8–10], it must be actively loaded onto, and unloaded from, DNA. In eukaryotes, PCNA is loaded onto DNA by the clamp loader ATPase RFC and CTF18-RFC and is actively unloaded from DNA by the ATAD5-RFC complex (ELG1-RFC in yeast) [11–14]. Dedicated clamp loaders and unloaders enable the cell to regulate when, where, and how much PCNA is loaded onto DNA through multiple different mechanisms. For two examples, the cell ensures cohesion establishment by co-localizing the CTF18-RFC clamp loader with stalled replication forks [15], and the cell signals for removal of PCNA by the ATAD5-RFC unloader by SUMOylating PCNA.

The stringent regulation of DNA replication and repair required for proper cell function places PCNA under immense selective pressure. Often, this pressure does not select for the “best” biophysical properties (e.g., the tightest binding afinities for PCNA’s partners) but rather maintains properties within an ideal “Goldilocks zone,” neither too strong nor too weak. Indeed, a landmark study by Fridman *et al*. demonstrated that increasing PCNA’s afinities for certain binding partners is more deleterious *in vivo* than completely abolishing those same interactions [16]. Just as PCNA’s binding afinity with its partners is finetuned in a Goldilocks zone, so too are the interactions that drive PCNA trimerization and ring formation. If the interaction between PCNA subunits were too tight, the energy converted from ATP by the clamp loader would be insuficient to pry open PCNA. On the other hand, if the interaction between two PCNA subunits were too weak, PCNA would spontaneously open and dissociate from DNA before its binding partners could perform their tasks.

Recently, our lab described the effects of a disease-associated PCNA variant C148S [17]. Patients who are homozygous for the PCNA-C148S allele display symptoms consistent with <u>P</u>CNA-<u>a</u>ssociated DNA repair <u>d</u>isorder (PARD). We showed that the PCNA C148S variant possesses a thermostability defect (ΔT_m_ = 10 K), affecting PCNA’s stability at physiological temperatures. We proposed that this stability defect results in spontaneous opening of PCNA and its passive unloading from DNA, driving the disease. To investigate the structural mechanisms by which the C148S substitution destabilizes PCNA, we determined the crystal structure of PCNA-C148S. However, there were no obvious differences between the variant and WT PCNA (overall 0.85 Å RMSD). Thus, it remained unclear how the mutation disrupts PCNA stability.

Here, we present extensive (~100 microseconds) classical molecular dynamics (MD) simulations (**Table 1**) and well-tempered metadynamics (WT-MetaD; **Table 2**) data that rationalize how the C148S substitution destabilizes PCNA. We predict that the C148S substitution interacts with water to destabilize an interfacial α-helix responsible for inter-subunit communication. By disrupting this α-helix, the C148S substitution destabilizes the inter-subunit interface and ultimately PCNA’s ring assembly.

**Table 1.** Equilibrium molecular dynamics simulations.

| **Equilibrium sampling** | **Replicates** | **Minimum replicate length (µs)** | **temperature (K)** |
|---|---|---|---|
| WT trimer | 3 | 2.5 | 315 |
| C148S trimer | 3 | 2.5 | 315 |
| WT dimer | 3 | 5 | 315 |
| C148S dimer | 3 | 5 | 315 |
| WT monomer | 3 | 6 | 315 |
| C148S monomer | 3 | 8 | 315 |

**Table 2.** Well-tempered metadynamics sampling.

| **WT-MetaD sampling** | **Replicates** | **Replicate length (µs)** | **temperature (K)** |
|---|---|---|---|
| WT trimer α-helicity=6 | 3 | 2.05-2.16 | 315 |
| C148S α-helicity=4 | 3 | 2.25-2.35 | 315 |
| C148S α-helicity=6 | 3 | 2.02-2.41 | 315 |
| WT dimer | 3 | 0.90-1.17 | 315 |
| C148S dimer | 3 | 0.81-1.06 | 315 |

## Results

### C148S can distort local α-helicity by hydrogen bonding with water molecules

To gain insight into how the C148S affects the conformational dynamics of PCNA, we performed equilibrium molecular dynamics simulations of WT and C148S at 315 K. We chose 315 K because we previously showed that the C148S variant is thermosensitive and the sensitivity is apparent at 315 K [17]. We considered three configurations: trimeric, dimeric, and monomeric PCNA (**Table 1**). We reasoned that trimeric PCNA simulations would provide insights into the most geometrically constrained and biologically relevant conformations, dimeric PCNA simulations would highlight the differences between an exposed and buried interface, and monomeric PCNA simulations would allow us to understand whether differences between WT and C148S arise within a monomer or if the differences require higher order assembly of PCNA.

Compared to WT PCNA, we find that the α-helix containing C148S has propensity to distort from idealized geometry. Specifically, the backbone hydrogen bond between F144 and C148S is transiently broken in all three oligomeric configurations (**Fig. 2A&B, Figs. 2S1-S4**). Although this conformational change is transient, it is fairly common, with trimer simulations spending ~15% of their time in the distorted conformation (compared to ~1.8% in WT simulations). When this occurs, the backbone hydrogen bond donors and acceptors of F144 and C148S are solvent exposed and create a hydrogen bonding cage with contributions from the C148S and T216 sidechains that can capture water molecules (**Fig. 2B, inset**). This cage is not formed by similar distortion of the WT α-helix, because the sidechain of C148 is a poor hydrogen bond donor and bulkiness of the thiol group tends to exclude water.

**Figure 1.**
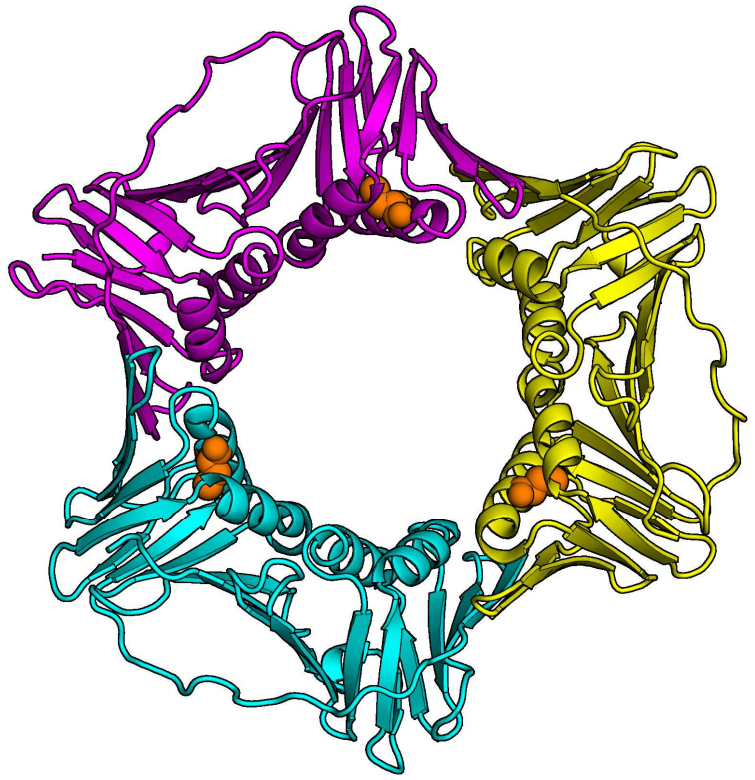
Human PCNA structure (PDB 1VYM). The three PCNA subunits are shown in magenta, yellow, and cyan, and the C148(S) position is shown as orange spheres, highlighting its proximity to the subunit interface.

**Figure 2.**
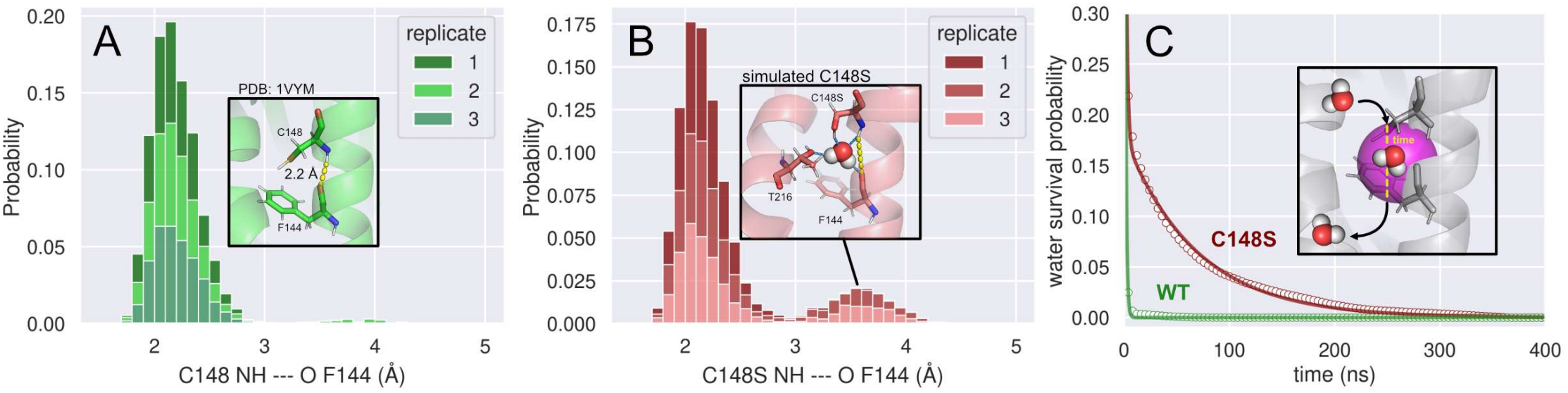
C148S destabilizes local secondary structure. (**A**) Normalized sampled probability distribution of the distance between F144 backbone oxygen and C148 backbone NH in trimeric WT PCNA simulations. (**A; inset**) The crystal structure of WT PCNA (PDB: 1VYM) shows a stable hydrogen bond (dashed yellow line) between the backbone of C148 and F144 (hydrogen atoms added to the crystal structure for direct comparison). (**B**) Normalized sampled probability distribution of the distance between F144 backbone O and C148S backbone NH in trimeric C148S PCNA simulations. (**B; inset**) A representative frame visualizing the stretched backbone distance (dashed yellow line) and hydrogen bond cage capturing a water molecule (hydrogen bonds shown by dashed blue lines). (**C**) Survival probability of a water molecule within a 5 Å radius sphere centered at the center of mass of C148(S) and F144 from trimeric WT and C148S simulations. (**C; inset**) The volume is shown as a magenta sphere, and the waters are depicted for schematic purposes.

To quantify how stably this cage can capture water molecules, we calculated the survival probability of water molecules entering a sphere with radius 5 Å centered at the center of mass of C148(S) and F144 (**Fig. 2C, Fig. 2S5**). We observe that when a water molecule enters this region in WT PCNA simulations, it is quickly excluded. When modeled to a single exponential decay, water is excluded from the WT site with time constant τ=1.08 ± 0.01 ns. However, we find that water molecules can survive in this region for upwards of 200 ns in C148S. To model these long lifetime events, the C148S survival probability must be fit to a double exponential. The time constants of the two phases of water exclusion from the C148S site are τ_1_=1.45 ± 0.03 ns and τ_2_=68.13 ± 0.74 ns. Because the fast time constant of C148S water exclusion roughly matches that of WT, we speculate that these two time constants model water exclusion when the α-helix is unperturbed and that the slow phase of C148S accounts for water molecules captured by the hydrogen bonding cage formed by the distorted α-helix. In support of this model, the amplitude of the slow phase accounts for ~18% of the data, roughly the amount of time we sample the distorted helix geometry. We note that our survival probability analysis likely represents a lower bound on total water occupancy lifetime, as it does not account for knock-on effects (i.e., when one water molecule is immediately replaced by another water molecule without restoring secondary structure).

### Residues flanking C148 are critical for inter-subunit communication but not for intra-subunit communication

Because the α-helix containing C148 lies at the inter-subunit interface, we hypothesized that residues within that α-helix are important for coordinating inter-subunit dynamics, but we were unsure whether they are also important for coordinating intra-subunit dynamics. To test the importance of this structural element for coordinating inter- and intra-subunit dynamics, we calculated linear mutual information from our molecular dynamics simulations across all three oligomeric states. From the linear mutual information networks, we calculated the betweenness centrality of the resultant graph. Betweenness quantifies the relative importance of residues for communicating dynamics within a protein assembly. Residues with higher betweenness are more important to information flow within the communication network than those with low betweenness (**Fig. 3A**); more details are provided in the methods section.

**Figure 3.**
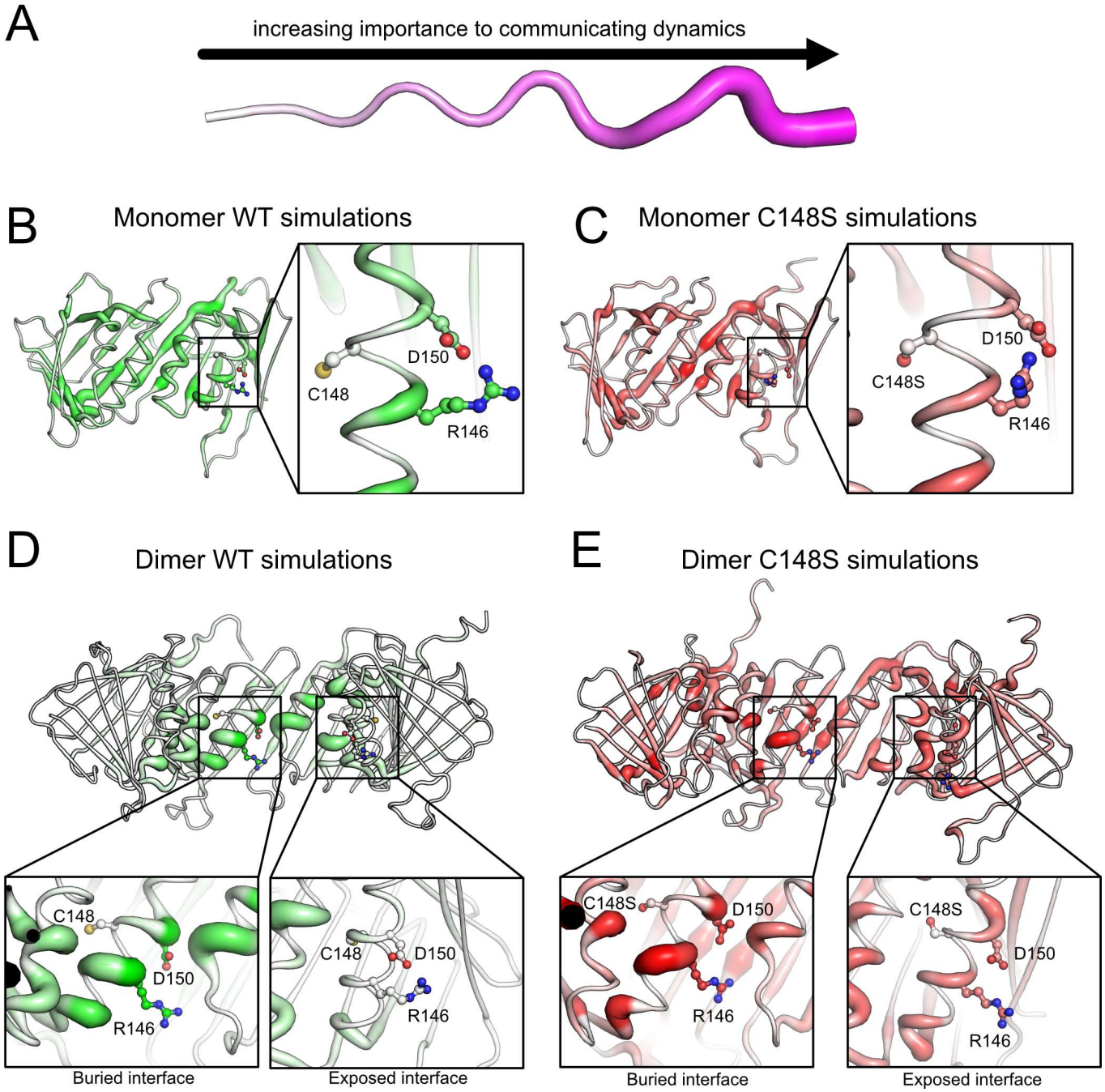
The residues flanking C148(S) are important for inter-, but not intra-subunit communication. **(A)** Schematic of betweenness representation. Residues with darker color and fatter tubes are more important for communicating dynamics. (**B and C**) Betweenness values calculated from our monomer WT and C148S PCNA simulations are shown in green-scale (**B**, WT) and red-scale (**C**, C148S). (**B and C; inset**) Close up views of the residues near C148 (**B**) and C148S (**C**) show that they are largely unimportant for coordinating intra-subunit communication. (**D and E**) Betweenness values calculated from our dimer WT (**D**) and C148S (**E**) PCNA simulations, shown with color-coding and tube thickness as before. (**D and E; inset**) Residues flanking C148 (**D**) and C148S (**E**) are important for mediating inter-subunit communication at the buried interface but are unimportant at the exposed interface.

In our monomer simulations of both WT and C148S PCNA, most communication is mediated by the central β-sheet, and little is mediated by the α-helix containing C148(S) (**Fig. 3B&C**). Similarly, we find that the solvent-exposed C148(S) α-helix in our dimer simulations also has low betweenness, which suggests that this α-helix does not play a significant role in intra-subunit communication. However, when inter-subunit contacts are present, we find that the residues flanking C148(S) (namely, R146 and D150) have very high betweenness, highlighting their importance for coordinating dynamics across the subunit interface (**Fig 3D&E, 3S1**). We note that our results are also consistent with D150 playing a critical role in maintaining PCNA trimer stability, as the conservative D150E mutation is known to disrupt trimerization [17–19].

From the analysis of our equilibrium simulations, we hypothesized that by destabilizing the local secondary structure of its α-helix, the C148S substitution can destabilize the PCNA trimer by affecting inter-subunit coordination. This could happen with at least two non-mutually exclusive mechanisms. First, by destabilizing the interfacial α-helix, the C148S mutation may strain the closed PCNA ring by biasing the angle between PCNA subunits towards values incompatible with closed-ring geometry. Second, by destabilizing the interfacial α-helix, the C148S mutation may directly destabilize the interface and promote spontaneous PCNA opening.

### C148S-induced α-helix destabilization does not affect equilibrium angle of PCNA dimer

Previous structural and computational studies of both bacterial and eukaryotic sliding clamps suggested that closed sliding clamp rings are under strain, because the subunits/dimers tend to flex outwards to geometries incompatible with closed ring geometry [20,21]. Thus, we hypothesized that by destabilizing the interfacial α-helix, the C148S mutation may add additional strain to the closed PCNA ring by further biasing the angle between PCNA subunits towards values incompatible with closed-ring geometry. To test this hypothesis, we performed well-tempered metadynamics (WT-MetaD) simulations of WT and C148S PCNA. WT-MetaD is a simulation technique that simultaneously enhances sampling and provides free energy estimates along reaction coordinates chosen by the researcher. We chose to employ two reaction coordinates: (1) the α-helicity of the helix containing C148S (as measured by the collective variable proposed in [22]), and (2) the flex angle between the PCNA subunits (**Fig. 4**). We chose a general reaction coordinate that describes overall α-helicity rather than using the distance between C148(S) and F144, as it remained possible that the disruption of the F144-C148S hydrogen bond (**Fig. 2B)** precedes further disruptions not captured in the timescale of our equilibrium simulations. The α-helicity collective variable ranges between 0 (no α-helical content) and 6 (ideal α-helix). We chose a flex angle that is ambivalent to whether the flexing is in-plane or out-of-plane because there are conflicting reports as to the importance of out-of-plane motions for clamp opening [21,23]; our flex angle uses less constraints on the system (requiring three centers-of-mass definitions instead of four that would be needed to define the out-of-plane dihedral) while allowing the dimers in our simulations to freely sample both in-plane and out-of-plane flexing. We performed three replicates for both WT and C148S starting from our equilibrium conformations; we report the average free-energy surface here in **Fig. 4**, and individual free-energy surfaces and the SEM of the average surfaces in **Figs. 4S1&2**.

**Figure 4.**
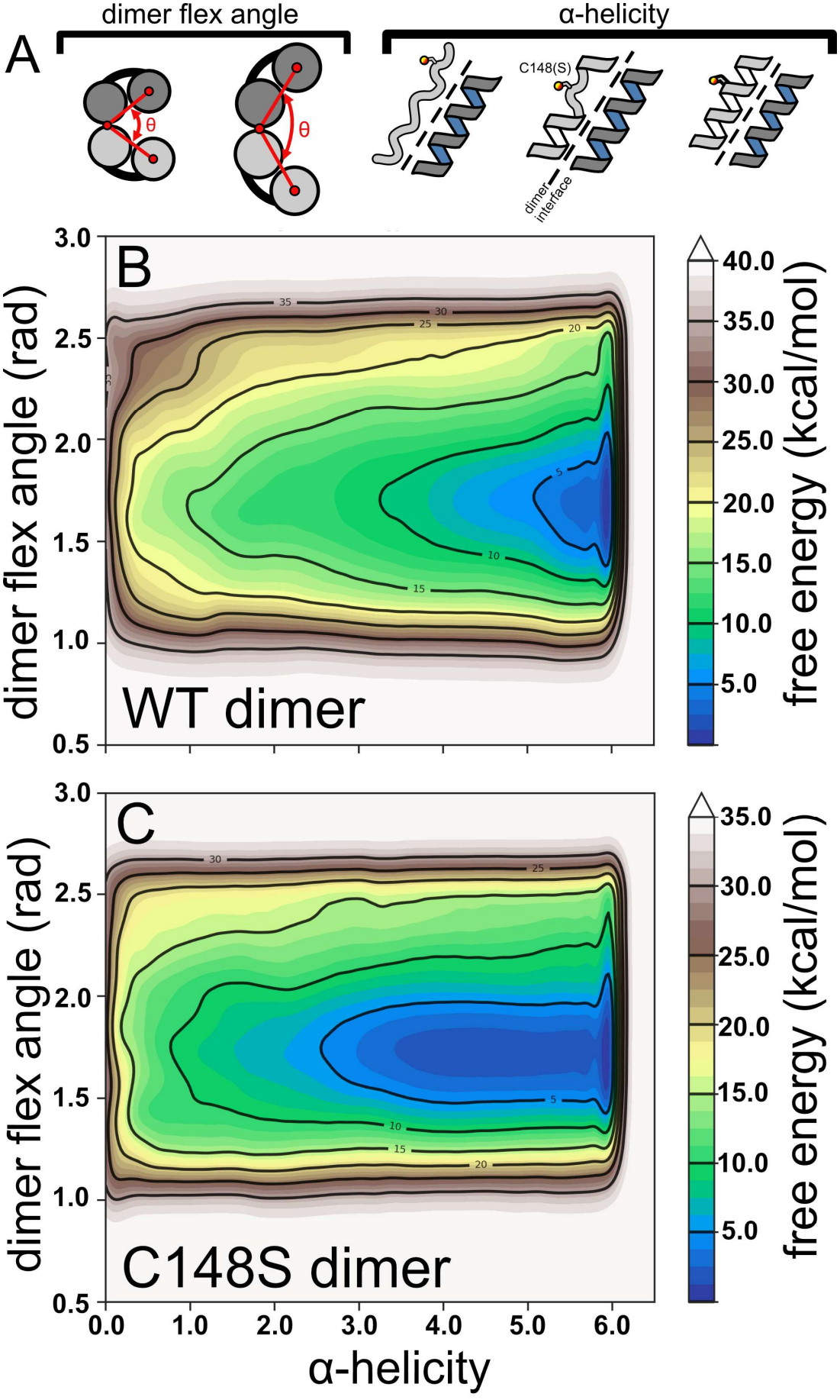
The C148S substitution disrupts helical stability but not the flex angle between PCNA subunits. (**A**) Schematics of collective variables. Two-dimensional free energy surfaces calculated from well-tempered metadynamics simulations of dimer WT PCNA (**B**), and dimer C148S PCNA (**C**). Each free energy surface is averaged from three independent starting conformations of PCNA, seeded from equilibrium simulations; the standard error of the mean and each replicate’s free energy surfaces are shown in **Figs. 4S1-2.**

Our calculated free-energy surfaces recapitulate the C148S helix destabilization observed in our equilibrium MD simulations by the appearance of a broad local energy minimum at *α-helicity* = ~4 that is not present in the WT free-energy surface. However, helix destabilization is not coupled with dimer angle flexing, suggesting the dimer angle can be constrained by other structural elements (likely β-sheet hydrogen bonding across the dimer interface). Thus, our data suggest that while the C148S substitution locally destabilizes its α-helix, this does not add additional strain to the PCNA ring by promoting inter-subunit angles that are incompatible with closed ring geometry.

### C148S-induced α-helix destabilization weakens the closed PCNA interface

Because it had been reported that the equivalent α-helix of *E. coli* β-clamp (α1″ in that complex) is distorted and that distortion is relieved by clamp opening [20], we next hypothesized that the α-helix distortion caused by the C148S substitution similarly destabilizes the closed PCNA interface, leading to spontaneous ring opening. To test this hypothesis, we performed additional WT-MetaD simulations that capture the opening of trimeric WT and C148S PCNA along two reaction coordinates: (1) inter-domain Cα contacts across a subunit interface and (2) inter-domain distance. Inter-domain distance captures large-scale opening of PCNA, while inter-domain Cα contacts capture localized destabilization of the interface that precedes complete opening. To test the contribution of the helix distortion found in our C148S equilibrium (**Fig. 2B**) and dimer metadynamics simulations (**Fig. 4C**), we further considered two simulation conditions for the C148S trimer: a stable helix (*α-helicity* harmonically restrained to 6) and a distorted helix (*α-helicity* harmonically restrained to 4).

We find that all three configurations (WT *α-helicity* = 6, C148S *α-helicity* = 6, and C148S *α-helicity* = 4) prefer the closed ring conformation (**Figs. 5, 5S1-3**). The identity of position 148 does not, by itself, substantially change the calculated free energy surface if the secondary structure of its α-helix is unperturbed (compare **Fig. 5A** and 5**B**). In either case, the closed ring conformation is preferred, inter-subunit contacts are maintained, and there is a large energetic barrier to ring opening. However, when the C148S α-helix is disrupted (**Fig. 5C**), we find that there is much lower energetic penalty for losing inter-subunit contacts and slight clamp opening; there is still a large energetic penalty for complete clamp opening. Thus, the C148S substitution may promote aberrant PCNA ring opening by first destabilizing local secondary structure which in turn disrupts inter-subunit contacts.

**Figure 5.**
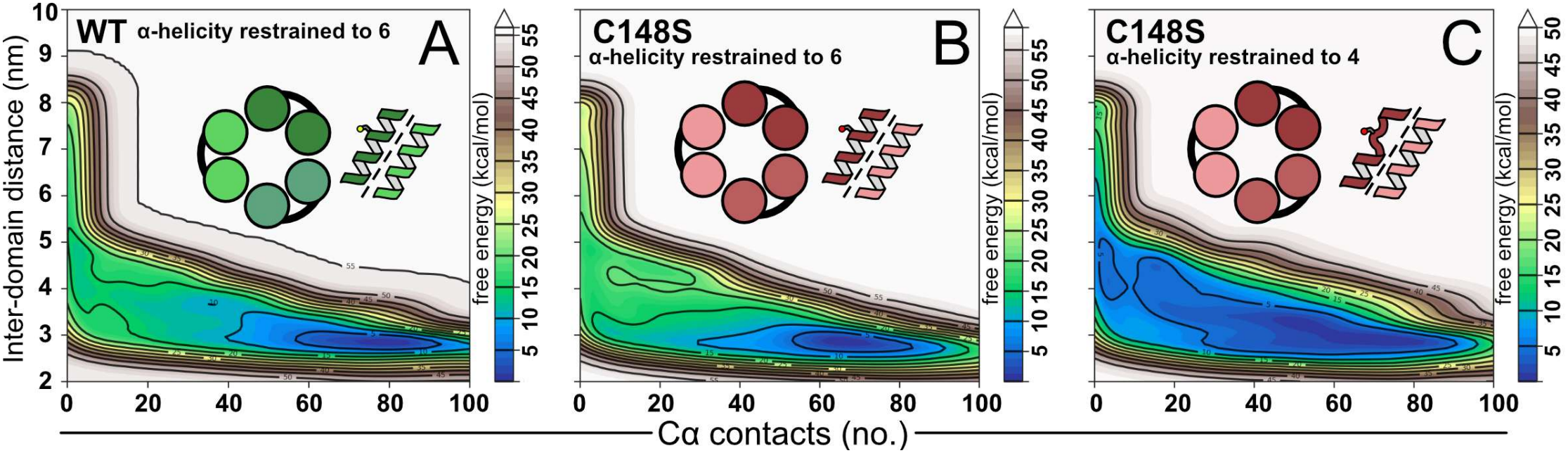
The C148S substitution can disrupt PCNA ring assembly. Two-dimensional free energy surfaces calculated from well-tempered metadynamics simulations of WT PCNA (**A**), C148S with maintained α-helix (**B**), and C148S with disrupted α-helix (**C**). Each free energy surface is averaged from three independent starting conformations of PCNA, seeded from equilibrium simulations; the standard error of the mean and each replicate’s free energy surface are shown in **Figs. 5S1-3**.

## Discussion

Patients harboring homozygous point mutations in PCNA can display a rare genetic disease, <u>P</u>CNA <u>A</u>ssociated DNA <u>R</u>epair <u>D</u>isorder (PARD) [17,24]. Our lab’s previous biophysical characterization of two such mutants showed that both variants display thermostability defects [17,25]. These findings suggest that understanding the molecular basis of PARD requires understanding PCNA’s dynamics. Molecular dynamics simulations are a powerful tool to gain insight into a protein’s dynamics, because they can predict conformational fluctuations and transient structural states that are inaccessible to traditional structural characterization methods. Thus, we simulated the dynamics of one such variant (C148S) and compared them against equivalent simulations of WT PCNA.

Our simulations shed light on the molecular mechanism by which the C148S substitution destabilizes the sliding clamp. We found that the C148S substitution transiently distorts the secondary structure of an α-helix that is critical for coordinating inter-subunit dynamics (**Figs. 2-4**), which in turn weakens the inter-subunit interface, leading to aberrant ring opening (**Fig. 5**). Importantly, the C148S α-helix nevertheless prefers WT-like secondary structure (as demonstrated by our equilibrium sampling in **Fig. 2B** and the global free energy minimum in **Fig. 4C**). Thus, probing this distortion is challenging experimentally and rationalizes why the defect was not observed in our crystal structure (PDB 8E84). The observed preference for WT-like helicity and ring structure was likely exacerbated by crystal packing forces and structure determination at cryogenic temperature, both of which favor the more compact and low energy WT-like conformation. Further, the preference for WT-like helicity and ring dynamics (compare **Fig. 5A** and **5B**) suggests that this variant can act generally like WT. This is consistent with our previous findings that C148S PCNA can be loaded onto DNA with WT-like eficiency and that, despite observed DNA repair defects, patients harboring this allele are viable [17].

More broadly, the rules governing the sliding clamp’s sequence-structure-dynamics relationship remain elusive. Sliding clamps across the domains of life have widely varying stability [26], despite strong structural similarity. For instance, although *Saccharomyces cerevisiae* PCNA and human PCNA crystal structures are nearly structurally identical (1.3 Å Cα RMSD between human [27] and yeast [28]), they only share a modest 35% sequence identity and human PCNA is less stable and more dynamic than its yeast homolog [29]. They also demonstrated that the backbone of α-helices lining the PCNA pore, including the helix containing C148, are more dynamic than the β-sheet core. Combined with our findings, this suggests that the C148S point mutation amplifies some basal instability already present within these α-helices. Towards the other end of the α-helix stability spectrum, the equivalent, WT α-helix in the *E. coli* sliding clamp is distorted in the closed ring conformation, and that distortion is relieved upon clamp opening; it was suggested that relieving the strain from α-helix distortion helps open the sliding clamp [20].

Thus, we propose that the stability of sliding clamp rings may be maintained within a “Goldilocks zone.” If the inter-subunit interface were too strong, clamp loaders and unloaders would not produce enough force to open the clamp. However, if the interface were too weak, the ring may spontaneously open and cause the sliding clamp to prematurely dissociate from DNA. There may be at least two competing forces dictating the balancing act: interfacial binding energy enforcing planarity of the closed ring, and the tendency for sliding clamp subunits to adopt right-handed helicity that matches the helicity of the clamp loader AAA+ modules. Previous molecular dynamics simulations of PCNA dimers were conflicted as to whether, without the constraint of the closed ring, PCNA subunits prefer to remain planar or match the right-handed helical pitch of the clamp loader [21,23]. However, these simulations were limited by sampling. In our several microseconds of sampling, PCNA dimers strongly prefer to adopt right-handed conformations (**Fig. 2S6**), suggesting that this tendency may also strain the ring and help clamp loaders/unloaders open the sliding clamp. Thus, with several factors at play it is difficult to predict the effects of a mutation in PCNA.

Lastly, we note that our previous biophysical characterization and current computational characterization presented here of the PCNA C148S variant suggests that researchers should use caution when designing cysteine-less or cysteine-lite versions of proteins. Indeed, detailed protein folding studies have revealed strong stability defects with mutations of cysteines to serine, particularly when buried [30–32]. While the cysteine to serine substitution is often considered conservative, it opens the potential for new hydrogen bond interactions with solvent that may be hard to predict even with state-of-the-art structure prediction algorithms (which often do not explicitly model solvent).

## Methods

### Simulation preparation

The initial structure of PCNA was taken from PDB 1VYM [27]. Because the crystal structure of C148S was nearly identical to WT PCNA but was unable to resolve the flexible Inter-<u>D</u>omain <u>C</u>onnecting <u>L</u>oop (IDCL), we chose to introduce the mutation into the WT crystal structure with the *swapaa* command in UCSF ChimeraX [33], which incorporates the most probable rotamer from the Dunbrack rotamer library [34]. The missing C-terminal residues were modeled using the Modeller server [35]. Structures were prepared for MD simulations with the tLEaP program from the AMBER suite [36]. The AMBER ff14SB force field was used to describe the protein [37]. PCNA trimers or dimers were centered in an octahedral periodic box of TIP3P water [38] with 20 Å padding to supply ample room for downstream simulations of PCNA opening or dimer flexing; PCNA monomers were centered in octahedral box with 14 Å padding. Neutralizing counterions and additional salt were added to reach 0.15 M NaCl [39] using Joung-Cheatham ion parameters [40]. CUFIX nonbonded corrections were applied to the entire system [41]. Topologies (.prmtop files) and initial coordinates (.inpcrd files) are included as supplemental files.

### Simulation protocol

Simulations were performed with OpenMM 8.X [42] on National Center for Supercomputing Applications Delta hosted at University of Illinois Urbana-Champaign or on UMass Chan High Performance Computing resources. A 1.0 nm nonbonded cutoff was applied, and long-range interactions were corrected with the Particle Mesh Ewald method [43]. Systems were energy minimized to remove steric clashes, followed by a 1 ns equilibration in the canonical (NVT) ensemble. Temperature was maintained at 315 K using Langevin dynamics (friction 1.0 ps^-1^). Next, systems were simulated in the isobaric-isothermal (NPT) ensemble using the Monte Carlo barostat to maintain 1 atmosphere of pressure (barostat interval 25 steps). Hydrogen mass repartitioning [44] was applied to allow for a 4-fs timestep to propagate the equations of motion. All systems were simulated in triplicate, and a summary of simulation trajectories can be found in **Table 1**. An example OpenMM input script is provided in the supplement.

### Distance and water survival probability analyses

To quantify the α-helix disruption, we calculated the distance between the backbone hydrogen of C148(S) and backbone oxygen of F144 after visual inspection of trajectories in VMD [45]. For dimer PCNA simulations, we considered each C148(S) site uniquely, as one site mediates inter-subunit contacts while the other site is solvent exposed. For trimer simulations, we considered each site to be indistinguishable and thus aggregated the data from all three sites. This segregating of dimer sites and aggregation of trimer sites was likewise employed to calculate survival probability of water near the mutation. To calculate the survival probability of water near the mutation, we used the *SurvivalProbability* function from the *waterdynamics* library implemented in MDAnalysis [46,47]. We defined the volume to be a sphere with radius 5 Å centered at the center of mass of C148(S) and F144. The data was fit to either a single exponential (functional form *A*_1_ exp 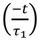 for WT trimer or double exponential (functional form *A*_1_ exp 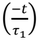 *A*_2_ exp 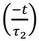 for C148S trimer using the *SciPy* Python library [48].

### Linear mutual information and betweenness calculations

To understand the relative importance of residues for coordinating motion within PCNA, we calculated the betweenness centrality of graphs constructed with linear mutual information using the *correlationplus* Python package [49]. Briefly, this method calculates the linear contribution to mutual information from the cross-correlation matrix of Cα fluctuations between residues. For our analyses, monomer and dimer PCNA simulations were aligned to minimize overall RMSD. To account for the symmetry of the closed trimer PCNA ring, we performed the linear mutual information calculation on pairs of dimers extracted from the overall trimer assembly. Thus, the mutual information calculation averages across all three interfaces from all three triplicate simulations rather than considering each interface to be unique.

From this calculation, one can construct a graph with nodes and edges connecting those nodes; nodes represent each residue of the protein, and the weights of the edges connecting nodes represent the amount of mutual information between the residues. Residues whose motions are tightly coupled are nodes connected with high edge weights, whereas weakly coupled residues are nodes connected with low edge weights. From this graph, one can calculate the optimal pathway connecting any pair of residues. The betweenness centrality of the graph is the proportion of times a node or edge is used in an optimal pathway in all such optimal pathways connecting all pairs of nodes. Thus, residues with high betweenness are more important for coordinating motions within a protein complex because they participate in more optimal paths.

### Well-tempered metadynamics protocol

To account for orthogonal degrees of freedom, the final frames from equilibrium sampling were used to seed starting configurations for independent replicates of well-tempered metadynamics (**Table 2**) [50]. Metadynamics simulations were run with OpenMM 8.X with PLUMED 2.9 *via* the *openmm-plumed* plugin [51,52]. All well-tempered metadynamics simulations were performed in triplicate and we report the average free energy surface; the resultant free energy surfaces of each replicate were shifted to set the minimum value to 0 prior to averaging. Each individual free energy surface, the standard error of the mean, sampling density, and Gaussian hill height decays are shown in **Figs. 4S1-2 and 5S1-3**.

For dimer metadynamics simulations, we chose two reaction coordinates (i.e. collective variables): the helicity of the α-helix containing C148(S) (*cv_helix)* and the angle between the dimer domains (*cv_angle). cv_helix* was defined as *ALPHARMSD RESIDUES = 404-414 (*residues 143-153 of the second monomer), and *cv_angle* was the angle between the centers of mass of the backbones of *com1*=(residues 133-251), *com2*=(residues 261-376), and *com3*=(residues 97-118, 427-444). Gaussian hills with height 1.0 kJ/mol that decayed with *biasfactor*=10 were deposited every 1,000 steps. To prevent unphysical states, upper and lower walls were used to constrain *cv_angle* to be between 1.047 and 2.500 rad. An example input script is provided in the supplement.

For trimer metadynamics simulations, we biased two collective variables: *cv_coord* and *cv_dist. Cv_coord* is the number of contacts between the Cα of residues 1-119 and 395-514 using the COORDINATION variable in PLUMED (*R*_*0*_=0.8 nm, *NL_CUTOFF*=1.2 nm, *NL_STRIDE*=100). *Cv_dist* is the distance between the centers of mass of the backbones of residues *com1*=(residues 1-119) and *com2=*(residues 395-514). Gaussian hills with height 1.0 kJ/mol that decayed with *biasfactor*=10 were deposited every 1,000 steps. To prevent unphysical collapse of the PCNA ring or interactions across the periodic boundary, upper and lower walls were used to constrain *cv_dist* between 2.2-9.5 nm, though our simulations never felt these walls. A harmonic restraint was applied to the *ALPHARMSD* collective variable (centered either at 6 or 4) of the α-helix containing C148(S) only at the opened interface. An example input script is provided in the supplement.

### Figures and plotting

Molecular representations were made with PyMOL [53]. Plots were generated with the *Seaborn* [54] and *Matplotlib* [55] Python libraries.

## Supporting information

MD topologies and example input scripts

## Data availability

Input files for molecular dynamics simulations are provided in the supplement for the bioRxiv preprint. All data associated with figures will be deposited in a permanent Zenodo repository upon peer review and final publication of the manuscript.

## Author contributions

J.P., J.M., and B.A.K. designed research. J.P. and K.B. performed research. J.P. and B.A.K. wrote manuscript and prepared figures. All authors edited and revised the manuscript.

## Acknowledgements

This work was funded by American Cancer Society award PF-22-114 to J.P., National Cancer Institute award F31CA254328 to J.M., and National Institute of General Medical Science awards R35GM156361, R01-GM127776, and R01-GM145943 to B.A.K. K.E.B. was supported by National Science Foundation award MCB-2342927 to B.A.K. This research used the Delta advanced computing and data resource which is supported by the NSF (award OAC 2005572) and the State of Illinois. Delta is a joint effort of the University of Illinois Urbana-Champaign and its National Center for Supercomputing Applications. The allocation for Delta was awarded through the NSF ACCESS mechanism, which is supported by NSF grants ACI-2138259, 2138286, 2138307, and 2138296. This research used computational resources provided by UMass Chan Scientific Computing for Innovation.

