## Supplemental Figures for "How a Disease-Associated Mutation Destabilizes the Sliding Clamp PCNA"

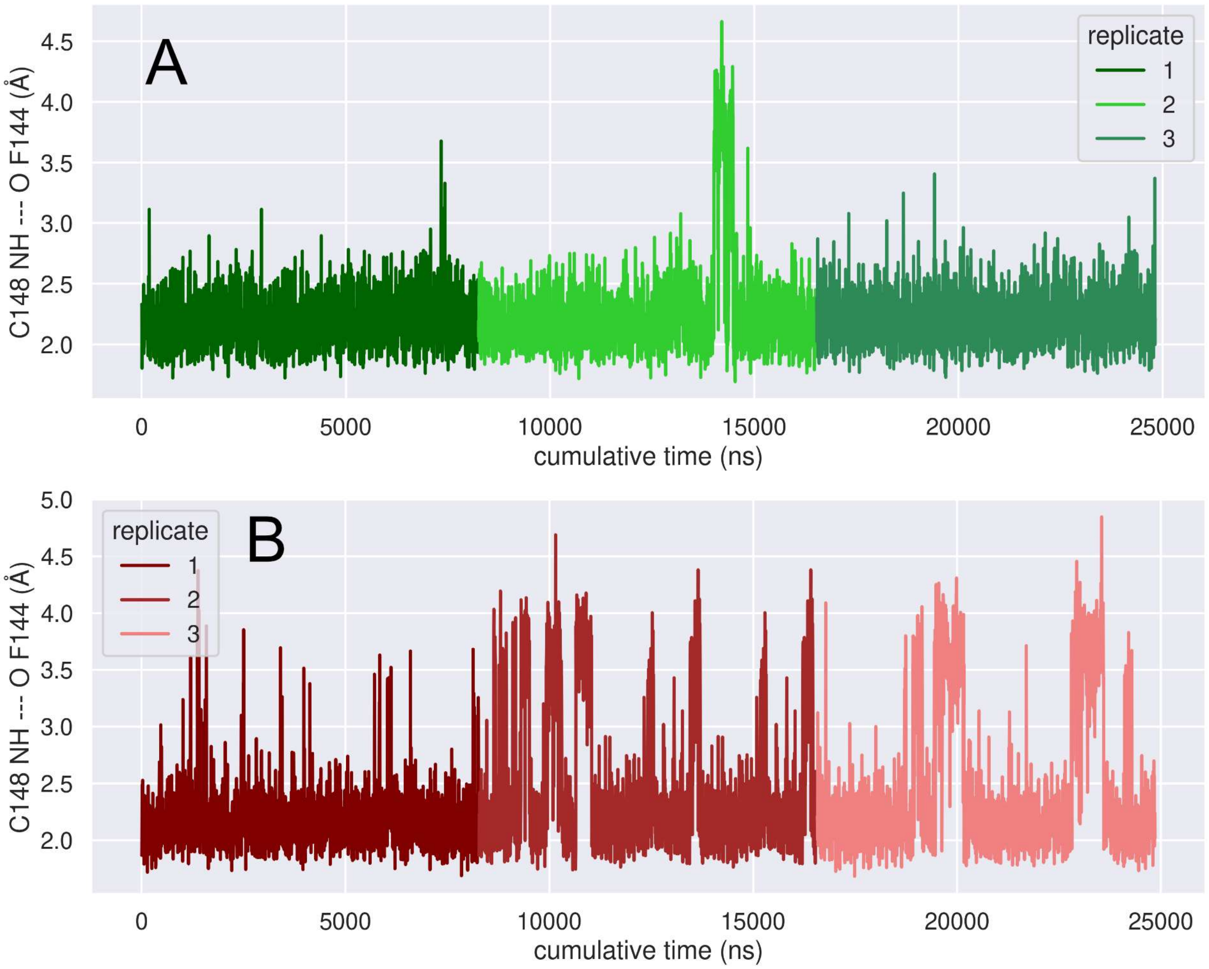


**Figure 2S1**. Backbone distance between F144 and C148 (**A**) or F144 and C148S (**B**) from trimer simulations. Each PCNA subunit is plotted in series within the same replicate.

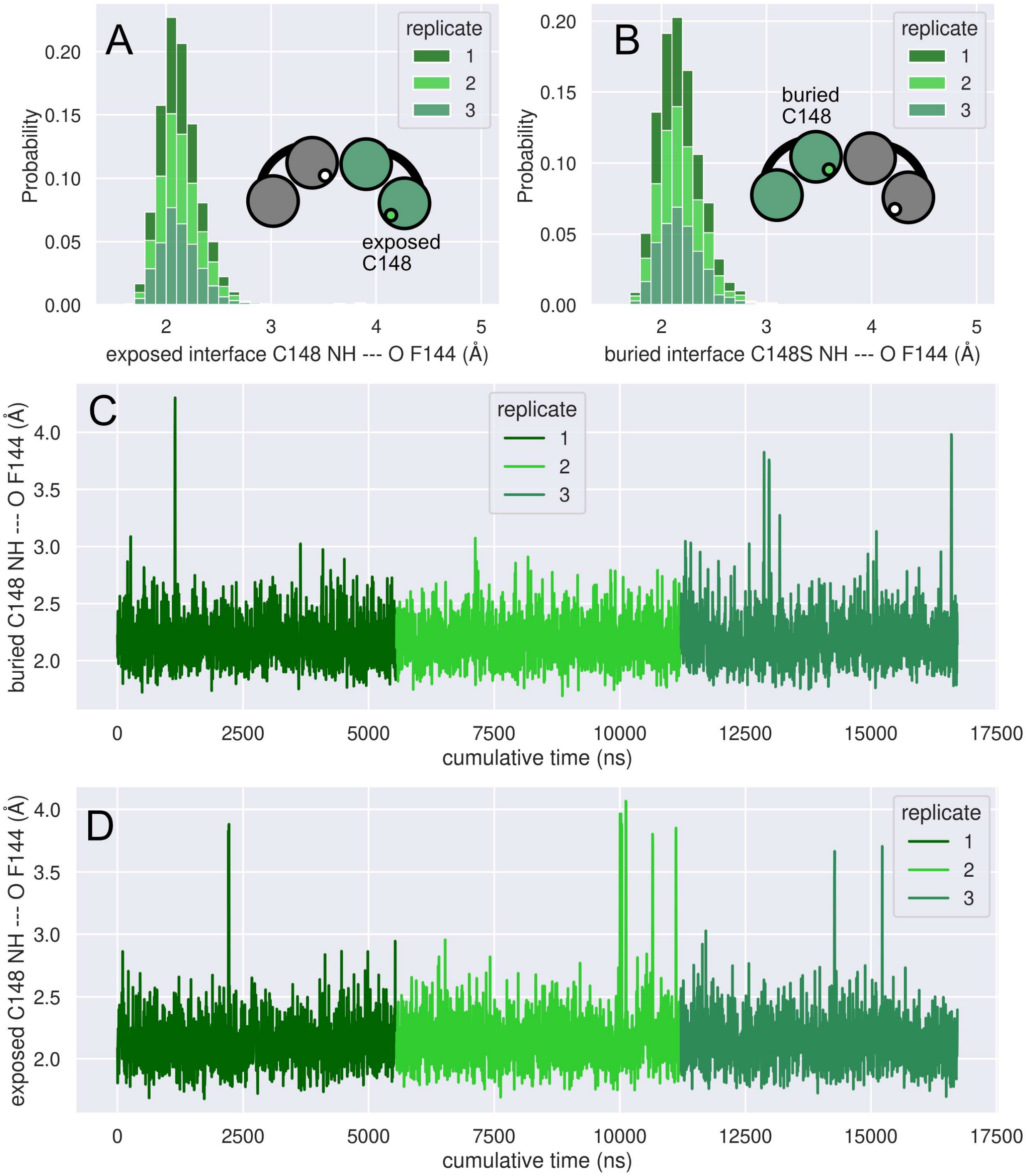


**Figure 2S2**. Backbone distances from WT dimer simulations. (**A**) Histogram of the backbone distance between F144 and C148 at the exposed interface. (**B**) Histogram of the backbone distance between F144 and C148 at the buried interface. Distance between F144 and C148 versus time at the buried interface (**C**) and exposed interface (**D**).

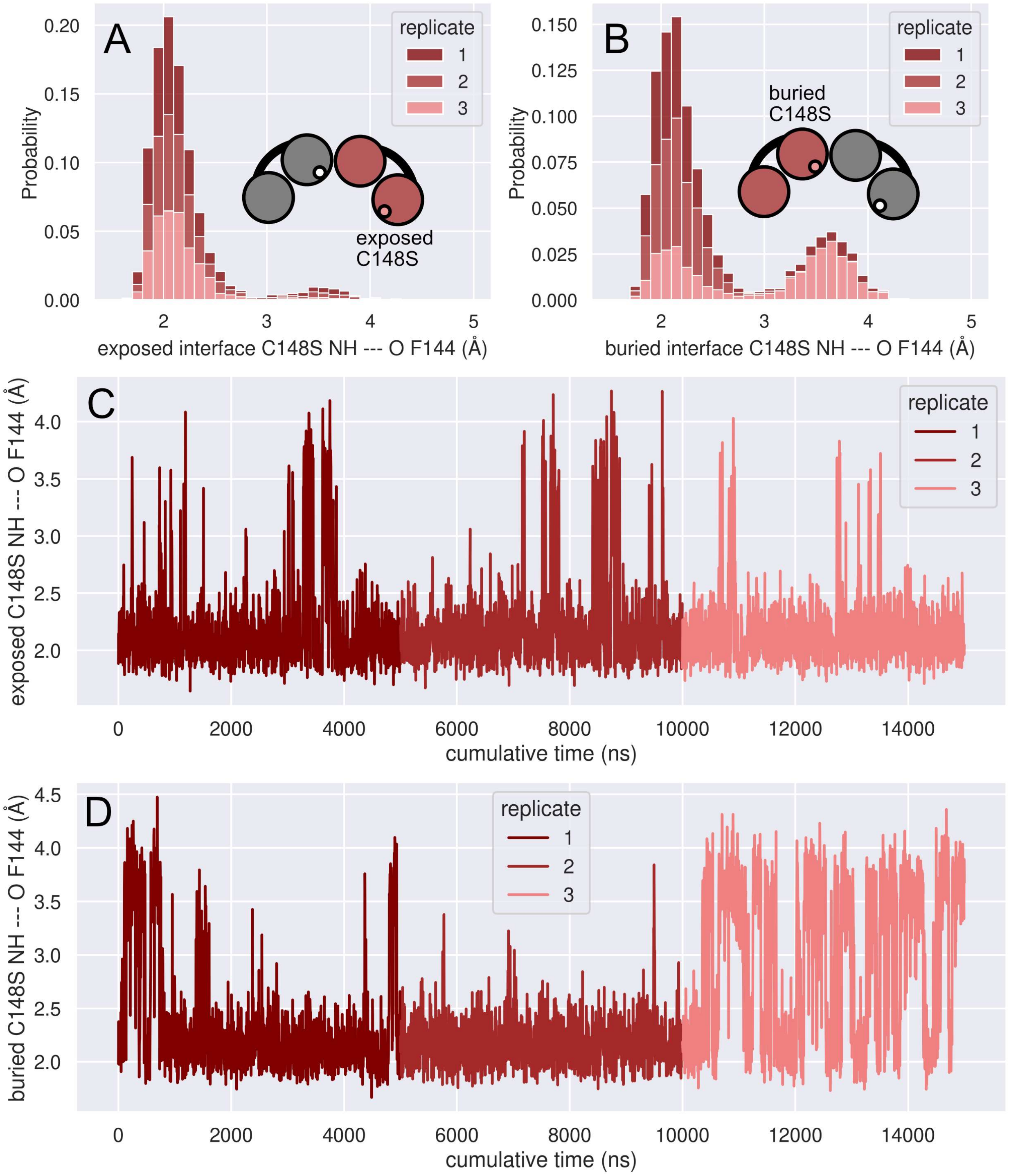


**Figure 2S3**. Backbone distances from C148S dimer simulations. (**A**) Histogram of the backbone distance between F144 and C148S at the exposed interface. (**B**) Histogram of the backbone distance between F144 and C148S at the buried interface. Distance between F144 and C148S versus time at the exposed interface (**C**) and buried interface (**D**).

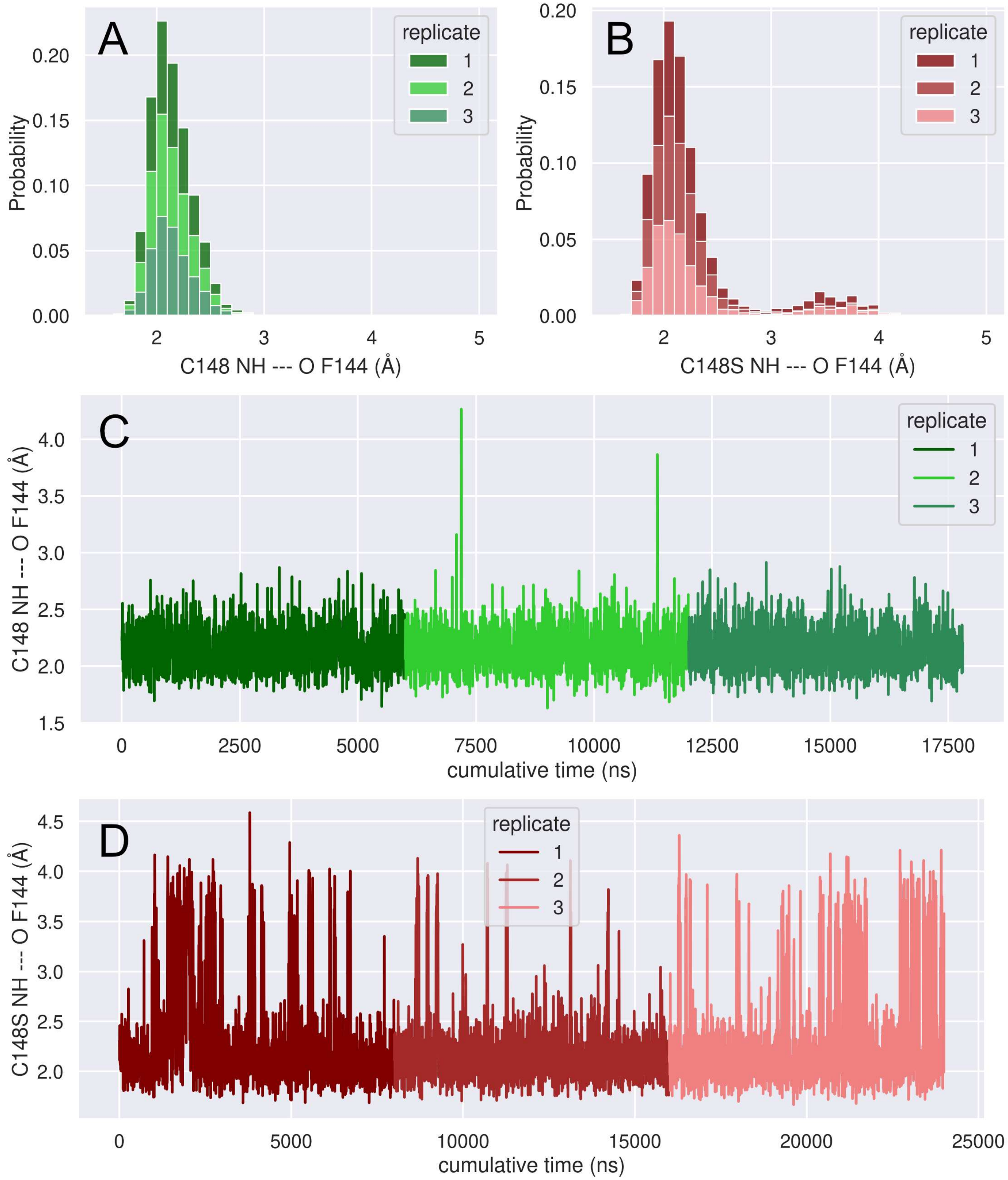


**Figure 2S4**. Backbone distances from WT and C148S monomer simulations. (**A**) Histogram of the backbone distance between F144 and C148. (**B**) Histogram of the backbone distance between F144 and C148S. Distance between F144 and C148 versus time (**C**) or between F144 and C148S (**D**).

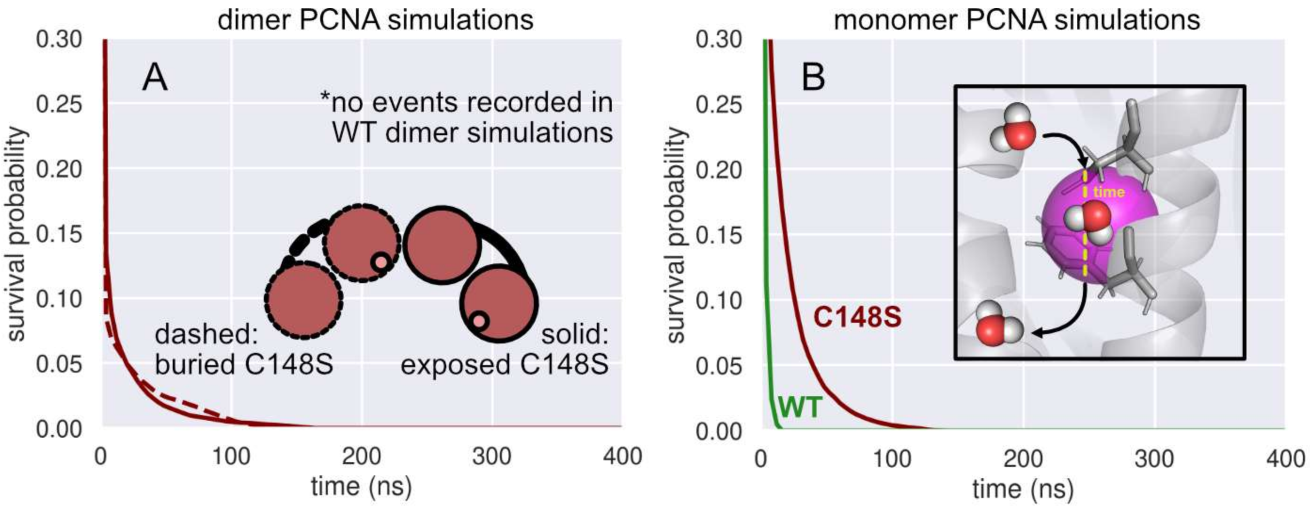


**Figure 2S5**. Water survival probabilities from dimer (**A**) and monomer (**B**) PCNA simulations.

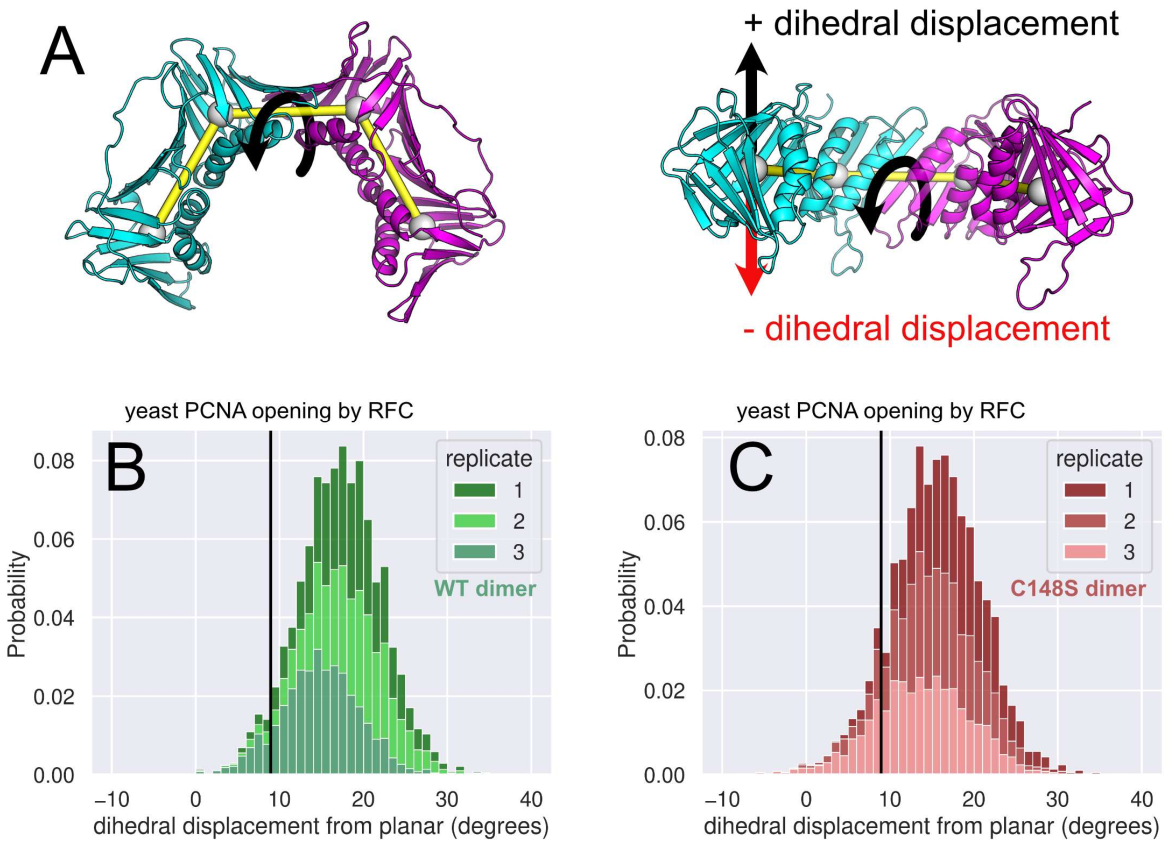


**Figure 2S6**. Dimeric PCNA adopts a right-handed conformation. (**A**) Helicity of dimer PCNA was calculated by the dihedral angle of the centers of mass of domains (gray spheres). The baseline helicity (~7.3 degrees) calculated from the planar crystal structure was subtracted, so that positive angles represent a right-handed helix and negative angles represent a left-handed helix. (**B**) Histogram of values sampled in the WT simulations. (**C**) Histogram of values sampled in the C148S simulations. Though the simulations were of human PCNA, the helical pitch of these domains when PCNA is opened by RFC was calculated from yeast RFC:PCNA structures (autoinhibited, PDB 7THJ; open, PDB 7TI8).

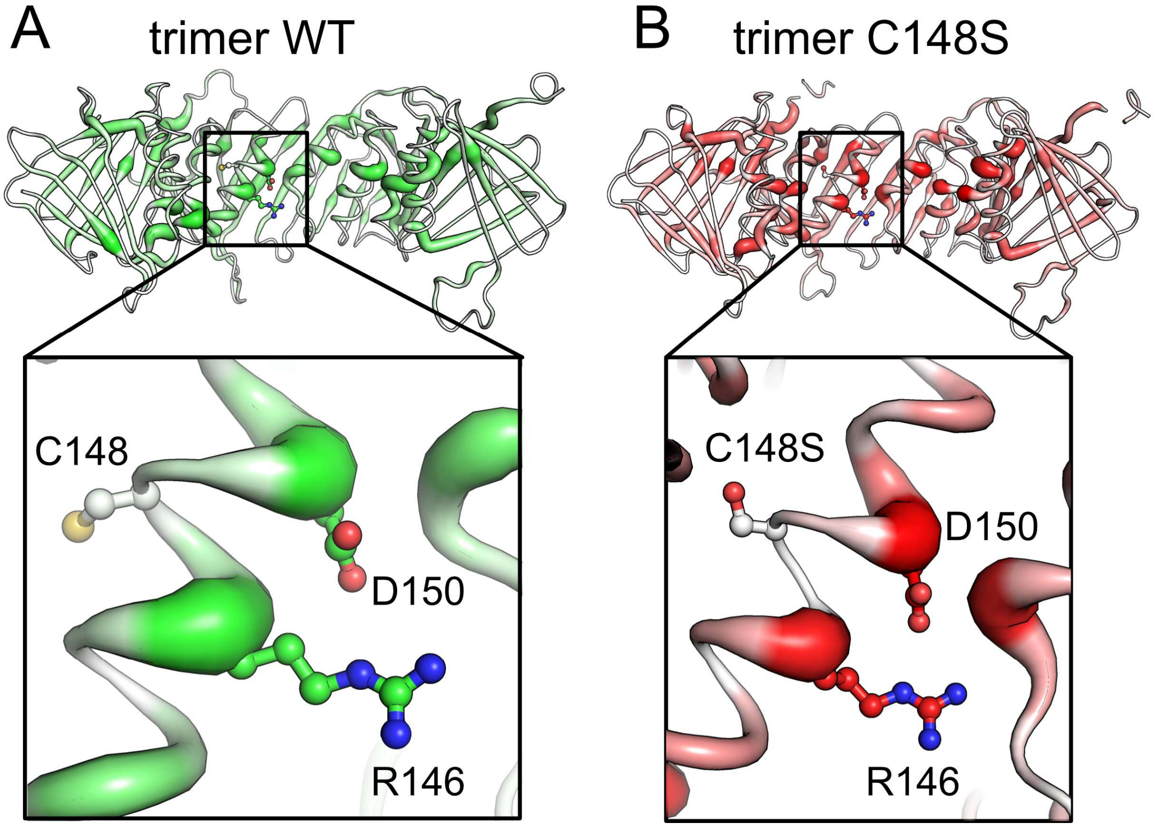


**Figure 3S1**. Betweenness calculated from (A) trimer WT simulations and (B) trimer C148S simulations show that D150 and R146 (residues flanking C148(S)) are important for coordinating dynamics across the inter-subunit interface.

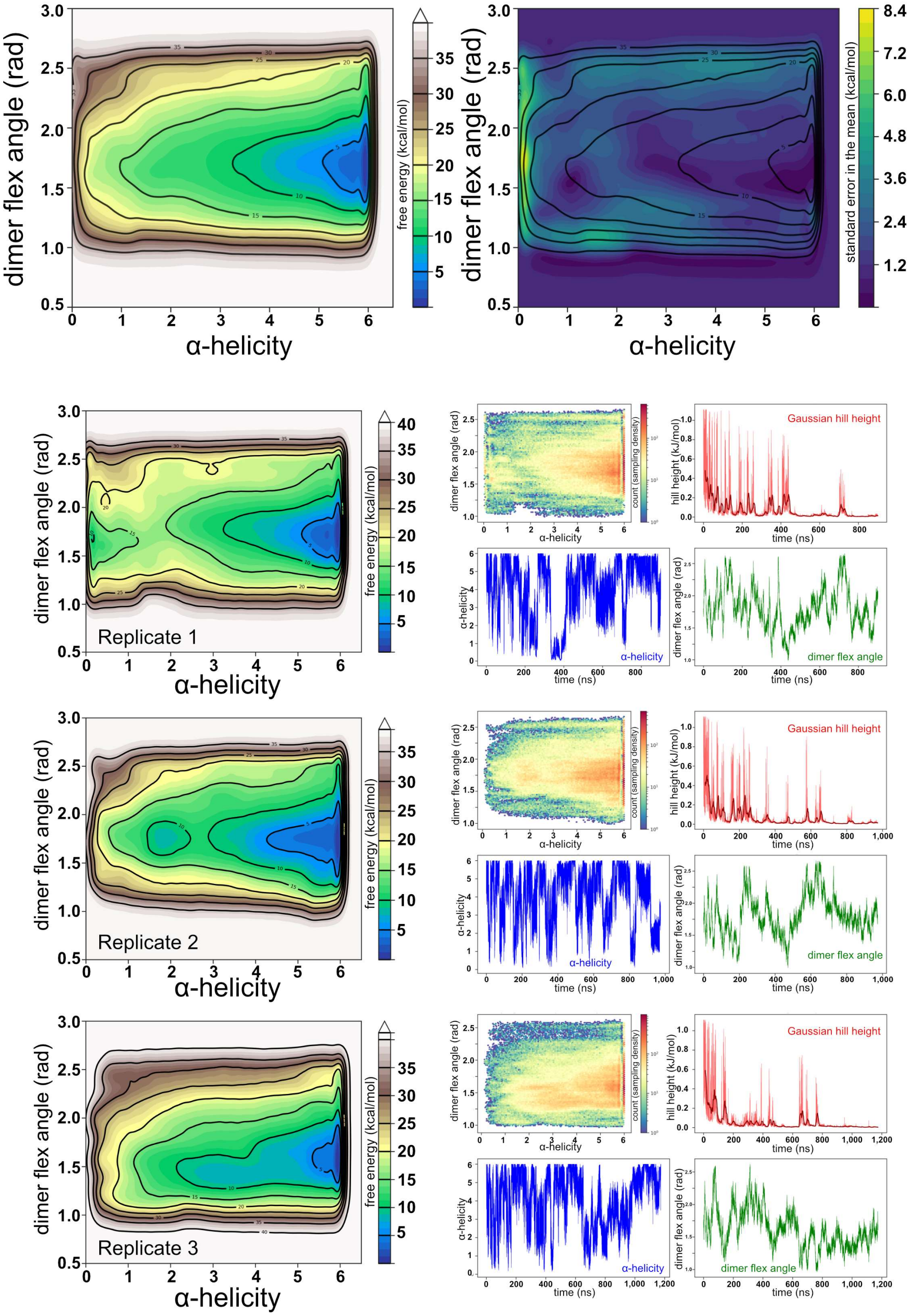


**Figure 4S1**. WT dimer metadynamics simulations. **Top**: Average free energy surface and standard error of the mean. Contours on the standard error are the contours of the free energy surface for easy visualization. **Bottom, lefthand column:** Each individual replicate free energy surface. **Bottom, righthand column:** Sampling density, Gaussian Hill heights (red), and individual collective variables (dimer flex angle: green; alpha-helicity: blue).

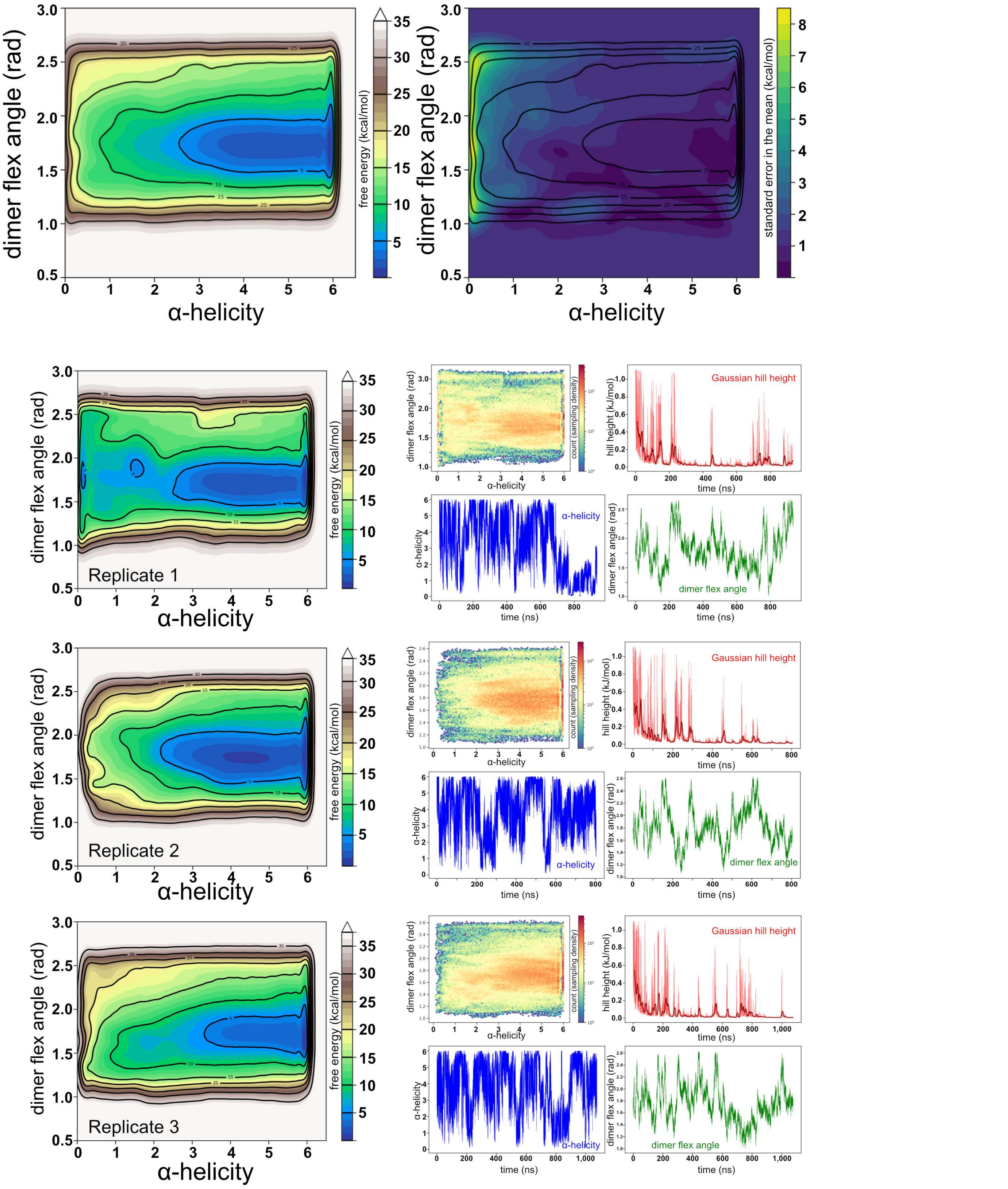


**Figure 4S2**. C148S dimer metadynamics simulations. **Top**: Average free energy surface and standard error of the mean. Contours on the standard error are the contours of the free energy surface for easy visualization. **Bottom, lefthand column:** Each individual replicate free energy surface. **Bottom, righthand column:** Sampling density, Gaussian Hill heights (red), and individual collective variables (dimer flex angle: green; alpha-helicity: blue).

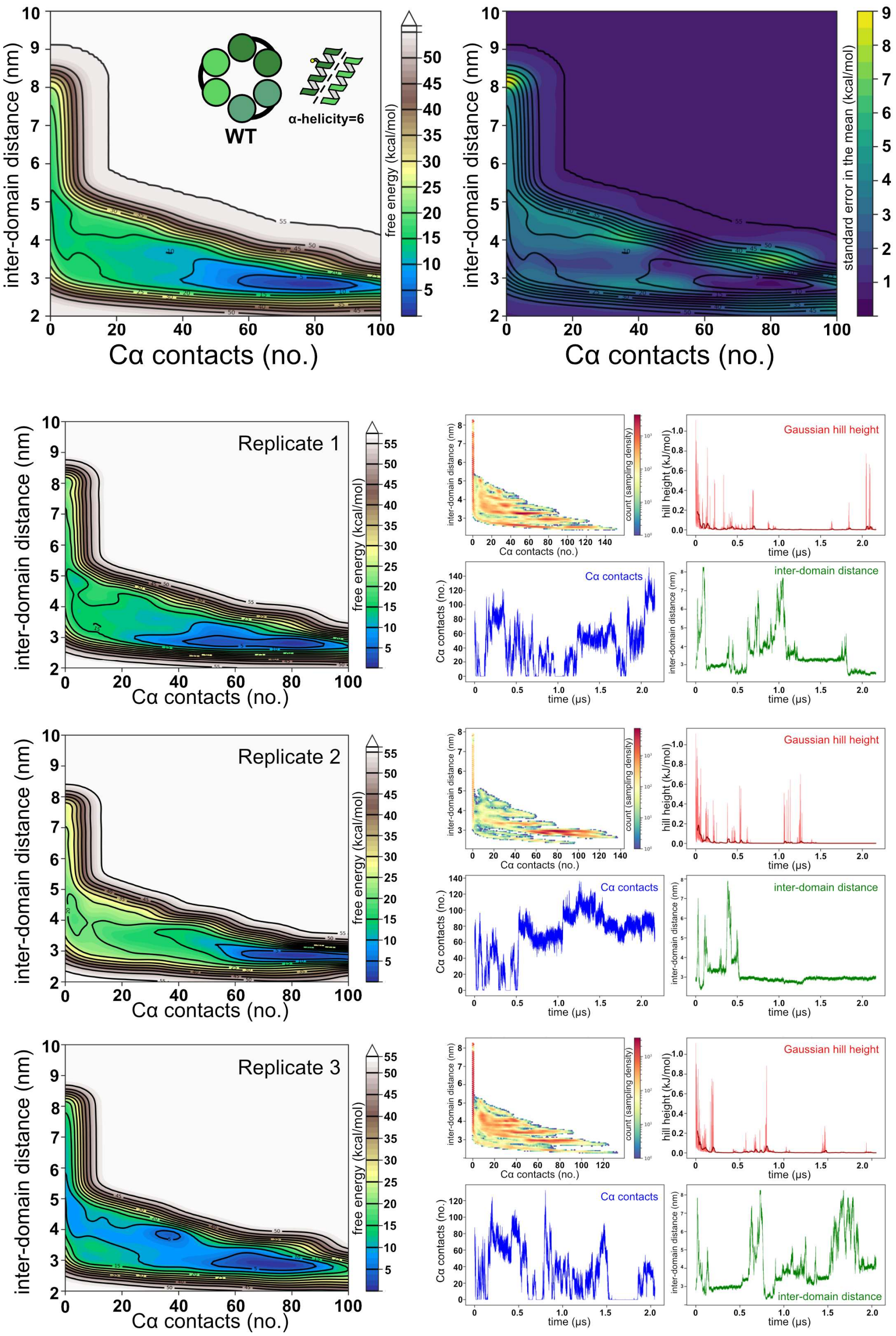


**Figure 5S1**. WT trimer metadynamics simulations (alpha_helicity=6). **Top**: Average free energy surface and standard error of the mean. **Bottom, lefthand column:** Each individual replicate free energy surface. **Bottom, righthand column:** Sampling density, Gaussian Hill heights (red), and individual collective variable traces (blue: coordination number, green: inter-subunit distance). In general we observe multiple opening events during each ~2 μs metadynamics simulation.

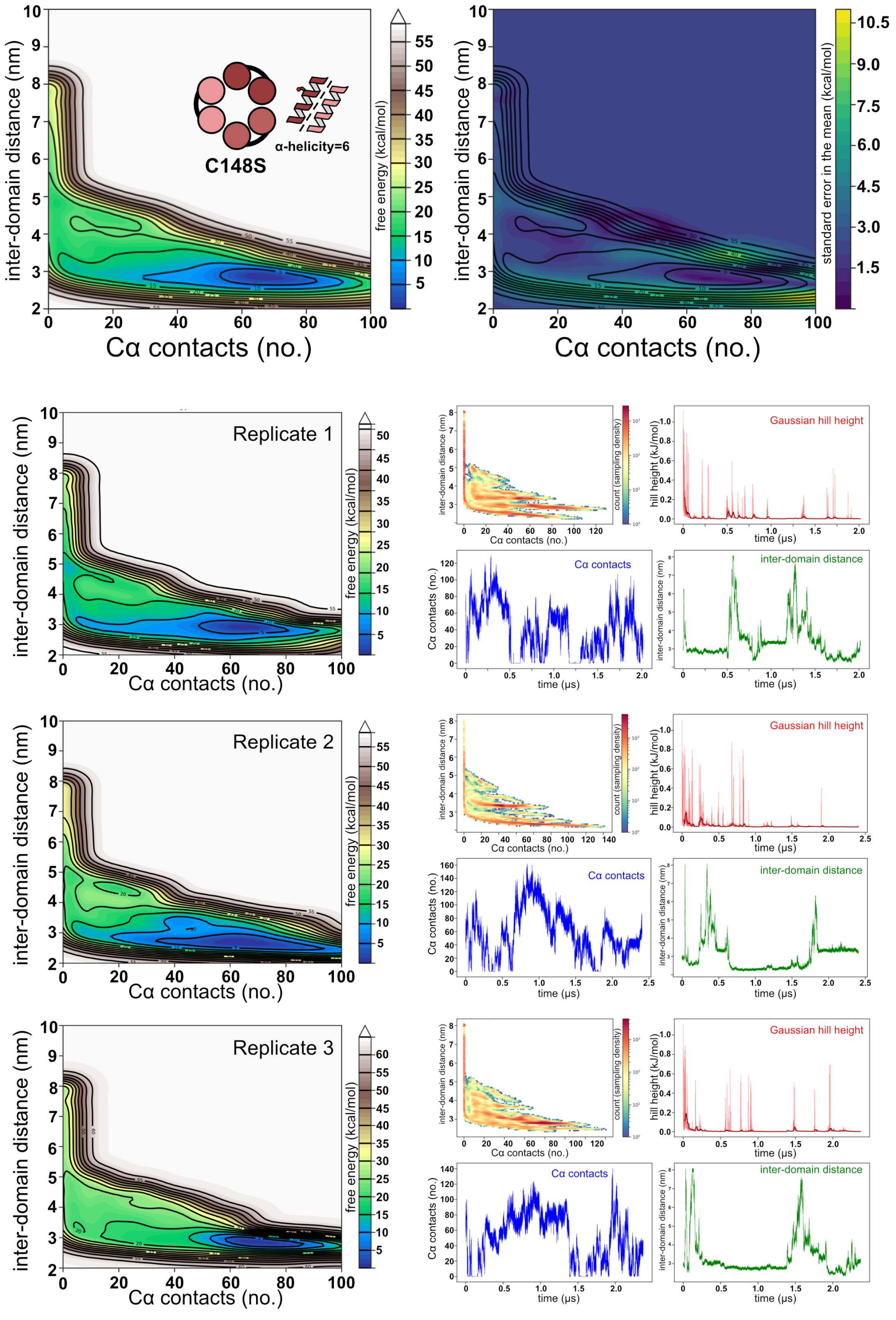


**Figure 5S2**. C148S trimer metadynamics simulations (alpha_helicity=6). **Top**: Average free energy surface and standard error of the mean. **Bottom, lefthand column:** Each individual replicate free energy surface. **Bottom, righthand column:** Sampling density, Gaussian Hill heights (red), and individual collective variable traces (blue coordination number, green: inter-subunit distance). In general we observe multiple opening events during each ~2 μs metadynamics simulation.

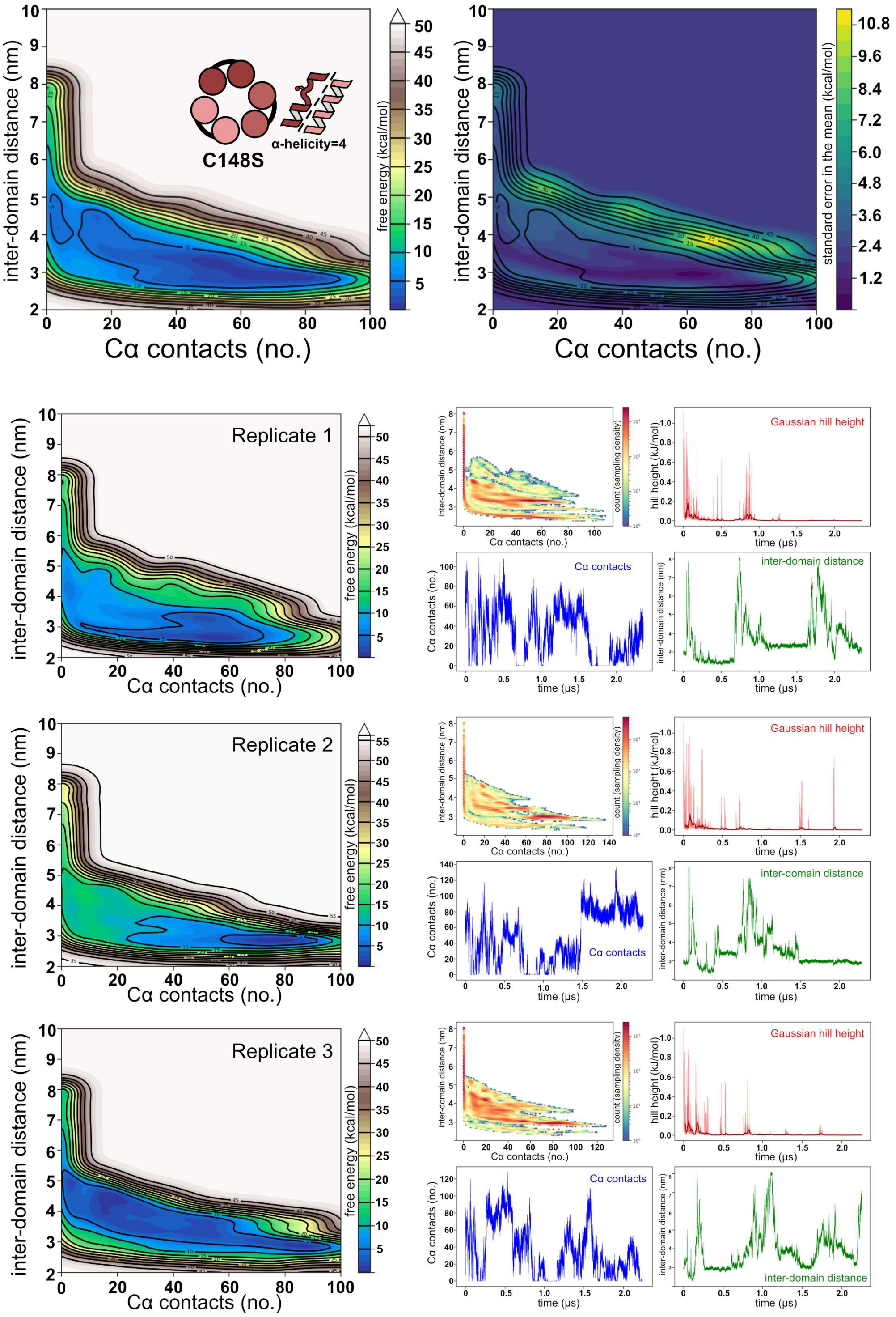


**Figure 5S3**. C148S trimer metadynamics simulations (alpha_helicity=4). **Top**: Average free energy surface and standard error of the mean. **Bottom, lefthand column:** Each individual replicate free energy surface. **Bottom, righthand column:** Sampling density, Gaussian Hill heights (red), and individual collective variable traces (blue: coordination number, green: inter-subunit distance). In general we observe multiple opening events during each ~2 μs metadynamics simulation.
